# geneSCOPE: gene Spatial Co-Occurrence of Pairwise Expression

**DOI:** 10.64898/2025.12.03.691993

**Authors:** Shicheng Zhang, Koichi Saeki, Hiroshi Haeno

## Abstract

Spatial transcriptomics captures neighborhood-dependent gene expression, but existing workflows do not always fully account for measurement scale and often treat space implicitly. We present geneSCOPE, a framework that integrates ecology-inspired statistics with network analysis. Molecules are binned on a grid whose width is chosen near the mode of the per-gene unit-invariant knee (UIK) distribution derived from Morisita’s *I*_*δ*_ –width curves. Pairwise adjacency-weighted spatial association is quantified with Lee’s *L*. We then assemble a spatial gene network and identify gene modules by consensus clustering. To identify cell–cell interactions between different cell types, high Lee’s *L* and low Pearson’s *r* is examined. Applied to human colorectal cancer (three Xenium sections) and a lymph node, geneSCOPE recovered spatial gene modules that map to microanatomy such as invasive margins, luminal epithelium, fibroblast-rich territories and germinal-center subdomains, and highlights intercellular neighborhood patterns at tumor–stroma interfaces between LGR5-marked stem-like tumor programs and C3-centered fibroblast/complement-associated niches. geneSCOPE thus provides a scalable, interpretable analytical foundation that generalizes across spatial omics and supports a wide range of applications.

## Introduction

Spatial context can substantially influence cellular phenotypes. *In situ* spatial profiling preserves the native spatial coordinates of molecules and cells within intact tissues, converting analyses of cell–cell interactions and neighborhood effects from inferred associations into directly measured features [1–3]. The value of this approach is evident in recent studies. For example, integrated single-nucleus and spatial transcriptomics of human pancreatic cancer delineated three reproducible multicellular neighborhoods, each comprising a characteristic mix of malignant, stromal, and immune cells with distinct local gene-expression programs [4]. At a larger scale, the mouse organogenesis spatiotemporal transcriptomic atlas (MOSTA) mapped continuous, fine-grained transcriptional gradients across mouse embryonic development, revealing that even canonical marker genes vary gradually rather than as step functions [5]. Similar spatial atlases are clarifying the organization of cardiovascular tissues and other disease contexts [6, 7]. Because physical proximity shapes phenotypes, spatial transcriptomic datasets often show locally correlated expression patterns—neighboring regions tend to share similar gene expression profiles [8]; analyses that ignore spatial coordinates risk missing these place-specific signals. Motivated by this spatial-first perspective, we introduce geneSCOPE, a framework that leverages geospatial information to map complex cellular neighborhoods and their neighborhood-specific gene modules in tissues.

For questions focused on short-range interactions, transcripts should be analyzed *in situ* without pre-aggregating by cell type. The analytical unit—whether an individual molecule, a grid bin, a spatial niche, or a tissue region—should match the biological question [1, 2, 4, 5, 7, 8]. Moreover, a spatially aware analysis must align with how measurements are made—the unit of observation, its spatial support, and its positional precision—because experimental design dictates achievable resolutions and appropriate statistics [9, 10]. Current platforms occupy complementary regimes of resolution and throughput: array-based technologies (e.g., 10x Genomics Visium) capture near whole transcriptome expression at multi-cell spots [1, 11], whereas single-molecule imaging (e.g., smFISH, MERFISH) achieves subcellular resolution but typically profiles targeted gene sets [12–15]; new high-definition *in situ* assays further increase molecule density, even in human tissues [16–18]. These trade-offs motivate a coordinated, variable-scale strategy rather than committing to a single spatial scale.

Several computational frameworks have been developed to jointly analyze gene expression and spatial position in transcriptomic data [19]. Giotto and Squidpy extend single-cell workflows with spatial adjacency graphs to enable domain detection and neighborhood-based clustering from tissue coordinates [20, 21]. More recent graph-based methods, including STAGATE and GraphST, leverage graph neural networks to integrate expression with spatial proximity, improving the delineation of tissue architectures and microenvironments [22, 23]. Probabilistic approaches such as cell2location combine single-cell reference signatures with spatial counts to estimate the abundance and co-localization of cell types across tissue sections [24]. Hotspot identifies spatially patterned genes and co-regulated gene modules, using neighborhood graphs rather than physical coordinates as explicit spatial weights, which may limit the capture of fine-scale physical adjacency in dense molecular maps [25]. Despite their utility, many approaches depend on specific platform resolutions or assume fixed neighborhoods, complicating generalization across datasets with differing measurement supports and molecule densities. Analytical tools should therefore flexibly adapt resolution and strategy to the characteristics of each dataset, balancing detail, noise, and computational cost.

Patterns of gene expression within tissues often resemble ecological distributions, where individual cells or transcripts occupy spatial niches and interact with their neighbors in structured ways. Concepts and analytical tools developed in ecology and geography—fields that study how organisms are distributed across space—can thus be applied to quantify biological spatial organization. Measures of global and local clustering (such as Moran’s *I* and LISA) detect non-random aggregation or segregation [26, 27], while hotspot statistics like Getis–Ord *G* identify regions of heightened or diminished activity, analogous to ecological “hot” and “cold” spots [28]. Bivariate association measures (e.g., Lee’s *L*) capture how two variables co-distribute across neighboring areas, and community indices such as Morisita–Horn describe the balance between mixing and segregation of populations and have shown prognostic value in oncology [29–31]. Together, these spatial-statistical approaches provide a common language for describing both ecological and molecular ecosystems, linking the organization of cells in tissue to the broader principles of spatial pattern formation.

In this paper, we present geneSCOPE—a spatially informed, variable-scale framework that (i) aligns to measurement scale, (ii) quantifies neighborhood co-distribution via spatially weighted gene–gene association, and (iii) organizes gene modules hierarchically to expose interfaces and cross-niche pathways. We demonstrate its utility and scalability on high-density human colorectal cancer and lymph node datasets.

## Methods

We developed a computational pipeline, geneSCOPE, to identify and cluster genes that exhibit spatially coordinated expression. The pipeline is implemented using R for data handling and orchestration, with compute-intensive kernels written in C++ and exposed through the Rcpp interface.

The workflow comprises five stages: (1) Grid-based spatial binning and normalization, (2) Spatial correlation analysis and robust inference, (3) Single-cell data integration, (4) Robust consensus network analysis, and (5) Network-derived hierarchy of spatial gene programs (Fig. 1). Each stage is described below.

**Figure 1.**
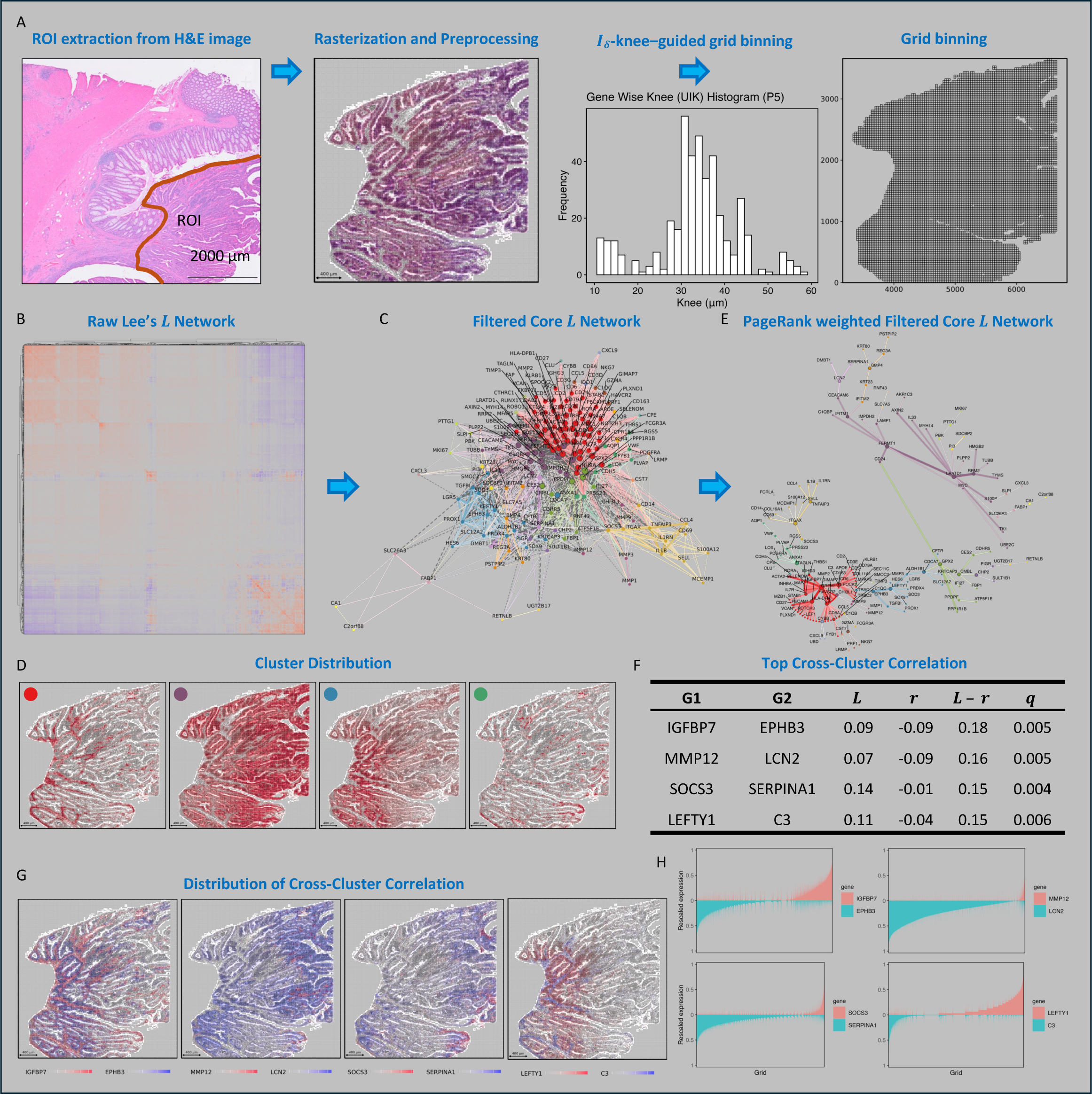
geneSCOPE workflow on CRC patient P5 (GSE280314) mapping spatial gene modules and neighborhood-specific gene–gene pairs. (A) A region of interest (ROI) at the colorectal cancer invasive front was defined using H&E staining, and transcripts within this ROI were aggregated into 30 μm square grid bins. The mode of histogram of per-gene UIK selects the working width. (B) Lee’s *L* values were computed for every gene pair on the normalized grid bin expression matrix, forming the initial spatial association network. (C) The spatial network was filtered by two criteria, retaining edges that (i) ranked in the top 5% by Lee’s *L* and (ii) remained significant after multiple-testing correction (*q* ≤ 0.05), resulting in a robust Lee’s *L* core network of gene co-occurrence. (D) Heatmaps show the spatial footprints of gene modules 1–4 (of 24 modules, derived from the core network). Each panel shows one module (1: red, 2: purple, 3: blue, 4: green), and color intensity encodes that module’s density per grid bin. (E) A PageRank-weighted dendro-network summarizes relationships among spatial gene modules. (F) A table highlights the top four gene–gene pairs exhibiting high Lee’s *L* but low Pearson’s *r* (*q* ≤ 0.05), indicating spatial co-occurrence that is enriched relative to their global single-cell co-expression. (G) Heatmaps depict the spatial distribution of each of these top *L* − *r* gene pairs across the grid bins of the tissue. (H) Mirror plots compare the distribution of each pair’s expression across the grid, highlighting patterns of local co-localization versus segregation. Abbreviations: CRC, colorectal cancer; ROI, region of interest; *L*, Lee’s *L* ; *r*, Pearson’s correlation.

### Grid-Based Spatial Binning and Normalization

To preserve native tissue geometry while reducing noise, we overlay a square grid on the region of interest (ROI) and assign each detected transcript to the nearest grid bin. The bin width is a tunable parameter anchored to the biological question and the assay’s spatial precision. When a priori length scales are unknown or multiple are plausible, we compute per-gene Morisita’s *I*_*δ*_ curves and estimate a UIK for each gene; the working width is chosen near the mode of the per-gene UIK distribution.

Morisita’s index *I*_*δ*_ is calculated as follows: for a gene in the *i*-th bin with counts *n*_*i*_ over *q* bins and total 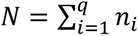,

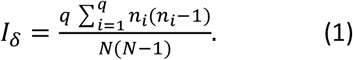

By convention, *I*_*δ*_ = 1 indicates a random (Poisson) distribution; *I*_*δ*_ > 1 indicates aggregation (clumping), and *I*_*δ*_ < 1 indicates uniform/regular dispersion. We compute per-gene *I*_*δ*_ across different grid bin widths (genes with *N* < 2 are excluded), then summarize across genes at each width. Very fine bins yield noisy or undefined values for low-abundance genes, whereas coarser bins stabilize estimates but blur local structure. To select an appropriate bin width, we locate the elbow of the *I*_*δ*_ –width curve using a unit-invariant knee (UIK) procedure: after unit-invariant rescaling of axes, the extremum-distance estimator chooses the point with the maximal perpendicular distance from the chord connecting the endpoints [32]. Then, the mode of the per-gene UIK (knee) distribution derived from Morisita’s *I*_*δ*_–width curves is selected as the bin width. In our datasets, this procedure typically selected 30–35 μm, balancing fine-scale detail and statistical stability. This binning improves signal-to-noise by pooling neighboring molecules while retaining neighborhood location via bin centroids.

Before aggregation, transcripts outside the ROI, technical controls, and putative background molecules (e.g., far from any nucleus when segmentation is available) are removed. Note that the ROI filter applies only to the coordinate-level spatial matrix; no out-of-ROI molecules enter the bin × gene matrix. After binning, each bin is annotated with polygon boundaries, centroid, covered tissue area, molecule count, and gene richness to enable overlays on histology. We then perform total-count normalization and compute gene-wise z-scores across bins to emphasize relative spatial patterning. The resulting bin × gene matrix is used in downstream spatial association analyses.

### Spatial Correlation Analysis and Robust Inference

We represent the grid as an undirected graph with the queen contiguity criterion [27], meaning that any two bins sharing a side or a corner are considered adjacent (each bin can have up to 8 neighbors including diagonally adjacent bins). Let *W* = {*w*_*ij*_} be the binary spatial weight matrix with *w*_*ii*_ = 0. Pairwise spatial association between genes *x* and *y* is quantified with Lee’s *L* statistic [33]:

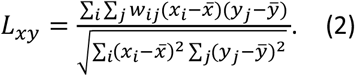

Here *x*_*i*_ and *y*_*j*_ are gene expression values in bins *i* and *j*, and *x̅*, *y̅* are means across bins. Positive *L* indicates spatial co-occurrence (high-with-high/low-with-low across adjacent bins), negative *L* indicates spatial avoidance, and values near zero indicate no spatial association.

To assess statistical significance of spatial patterns, we employ a combination of analytical and permutation-based methods. We use spatially constrained block permutations (preserving macro-architecture) to generate the null for *L*. Repeating this procedure 1,000 times yields a null distribution of Lee’s *L* values for each pair, against which we compare the observed *L* to compute an empirical p-value. All resulting p-values are adjusted for multiple testing to ensure robust inference. By default, we control the false discovery rate (FDR) using the Benjamini-Hochberg procedure, given the large number of gene pairs tested [34]. For all module discovery analyses presented in this study, we constructed a core Lee’s *L* network by retaining edges with positive *L* values that passed a FDR threshold of *q* ≤ 0.05 and then further restricting to approximately the top 5% of edge weights for CRC samples and the top 0.1% for lymph node (LN) after applying a *log*(1 + *L*) transformation.

### Single-Cell Data Integration Strategies

The geneSCOPE pipeline integrates single-cell (or single-nucleus) RNA-seq as either (i) a spatially matched reference that includes only cells within the ROI, aligning cell profiles to spatial bins or (ii) a comprehensive reference that includes all profiled cells. In both modes, single-cell counts are total-count normalized, placing single-cell expression on a comparable scale to the grid data and accounting for sequencing depth differences. The resulting single-cell matrix can then be directly compared or integrated with the spatial bin-level data.

We primarily use single-cell data as a reference to distinguish spatially driven gene relationships from general co-expression. For each gene pair, we compute the Pearson correlation across all single cells to get a baseline co-expression without spatial context, and we contrast this with the spatial association (Lee’s *L*) from the tissue. A gene pair with a high Lee’s *L* but low single-cell correlation indicates context-specific co-expression driven by spatial proximity. Conversely, if two genes are strongly correlated across single cells but not spatially co-localized, their co-expression likely reflects a cell-intrinsic program rather than a spatial interaction. By comparing spatial statistics with single-cell correlations, geneSCOPE can isolate neighborhood-specific co-enrichment from broader transcriptional programs.

### Robust Consensus Network Analysis

Significant spatial gene-gene correlations are assembled into a gene network, where nodes are genes and edges are weighted by their spatial correlation from calculated Lee’s *L*. To obtain robust modules, we run the Leiden community detection algorithm on this core gene correlation network 1,000 times with small random perturbations [35–37]. From these runs, we construct a consensus matrix indicating how often each gene pair co-clusters across runs and then perform community detection on this matrix to derive the final modules. For all analyses presented here, we define consensus modules used throughout the study as groups of genes that co-cluster in at least 95% of runs, require modules to contain at least two genes, and discard isolated or unstable nodes. This yields robust gene modules that are reproducible across different clustering runs.

To assess the coherence and distinctness of modules, gene–gene Jaccard analysis is performed. For each gene, we binarized its spatial footprint on the selected grid by marking bins as present if its expression exceeds a threshold (z-score > 0). The gene–gene Jaccard similarity matrix is computed as follows: 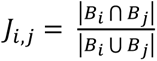. Using the final Leiden assignments, we summarized for each gene the difference between its mean Jaccard similarity to genes within the same module and to genes outside it with a paired Wilcoxon signed-rank test. At the module level, we binarized module footprints by thresholding per-bin module scores (z > 0) and computed a module–module Jaccard matrix to visualize spatial relationships with a heatmap.

Finally, we integrate the network results back onto the tissue image by visualizing each module’s activity *in situ*. For each gene module, we calculate a module expression score per bin and plot these scores on the tissue image. This generates a spatial map of each module’s “territory,” often revealing boundary or interface regions where a gene expression program is spatially coordinated. Such spatial mapping of modules helps validate that the network-derived gene communities correspond to meaningful anatomical or microenvironmental structures in the tissue.

### Hierarchical Analysis of Spatial Gene Modules with a PageRank-Weighted Core Network

We summarized relationships among spatial gene modules without changing module memberships. We treated the consensus gene network as a weighted graph whose edges encode spatial association strength (Lee’s *L*) between pairs of genes. We computed PageRank centrality scores using igraph package for each gene on this network (using a standard damping factor of 0.85) and used these scores to reweight edge strengths [35].

From this reweighted network, we extract a tree-like backbone using minimum spanning trees (MSTs) [35]. Within each module, we compute an MST that retains only the strongest edges required to keep all genes in that module connected. We then collapse each module into a single node, aggregate inter-module edge weights across all gene–gene connections between module pairs, and compute a second MST on this module-level graph. This two-stage MST procedure yields a tree that orders modules and defines their adjacency in terms of the highest-weight connections that are necessary for connectivity.

For visualization, we embedded this backbone using a tree or radial layout and overlay a small number of the highest-weight non-tree edges to preserve prominent alternative connections while keeping the graph sparse. In these layouts, modules joined by short paths of high-weight edges appear adjacent and are interpreted as closely linked spatial programs, whereas modules that are separated by longer paths correspond to programs that are only weakly coupled in the network. Because each module is defined by a distinct spatial footprint on the tissue, adjacency in this backbone often coincides with physical interfaces or gradual changes between module-enriched regions. This representation is based on symmetric spatial associations, does not encode temporal or causal direction, and is intended as an intuitive visualization that can help generate hypotheses about spatial relationships among gene modules.

### Data & Code Availability

The geneSCOPE source code and all scripts used to reproduce the results are available at https://github.com/CoooRossa/geneSCOPE under the MIT license. The Xenium human CRC dataset was generated from ref. [18], and the original data can be accessed from GEO under accession GSE280314. The Xenium human lymph node dataset was generated from ref. [38], and the original data can be accessed at https://www.10xgenomics.com/datasets/preview-data-xenium-prime-gene-expression.

## Results

### Overview of the geneSCOPE Workflow on a CRC Invasive-Front ROI

We first applied geneSCOPE to a human colorectal cancer (CRC) tissue section (10x Genomics Xenium dataset GSE280314, patient P5) to illustrate the end-to-end workflow (Fig. 1). A region of interest (ROI) centered on the invasive tumor front—characterized by densely packed nuclei and disorganized epithelium—was defined on H&E. Using the mode of the per-gene UIK distribution from Morisita’s *I*_*δ*_-width curves (Fig. 1A), we chose a 30 μm grid as the spatial analysis resolution for transcript binning.

We computed Lee’s *L* for all gene–gene pairs (Fig. 1B). Edges significant after FDR correction (*q* ≤ 0.05) were retained, and consensus community detection identified 24 modules of genes with similar spatial expression patterns (Fig. 1C). Because *L* weights spatial adjacency, each gene module corresponded to a distinct tissue domain at the 30 μm analysis scale, as confirmed by overlaying module expression scores onto the tissue image (Fig. 1D). To visualize inter-module relationships, we constructed a PageRank-weighted random-walk core from the L-weighted gene network and summarized inter-module proximity with a two-stage MST (Fig. 1E). Finally, we contrasted *in situ* spatial association (Lee’s *L*) with non-spatial co-expression from matched single-cell RNA-seq (Pearson’s *r*), prioritizing pairs with high *L* at low *r* (Fig. 1F–H). At the invasive margin, for example, C3–LEFTY1 co-localized within the same neighborhoods, and EPHB3–IGFBP7 appeared in proximity across epithelial–stromal interfaces—patterns consistent with context-dependent interactions. Applying geneSCOPE to two additional CRC samples yielded consistent results, demonstrating generalizability and utility (Fig. S1, S2).

### Mapping Canonical Niches in Human Lymph Node Using geneSCOPE

Using the same workflow, we next analyzed an FFPE human lymph node section profiled with the 10x Genomics Xenium 5K Pan-Tissue & Pathways gene panel (4644 genes in this case) [38]. A composite ROI was selected to encompass the H&E-defined light and dark zones of a germinal center (GC) along with adjacent lymph node regions. We performed 30 μm grid binning, normalization, Lee’s *L* computation, spatial network construction, and consensus clustering exactly as in the CRC analysis (Fig. 2). The resulting gene modules exhibited spatial footprints corresponding to known GC substructures and surrounding functional compartments, indicating that high-density molecule coordinates can partition tissue regions without prior cell-type annotation.

**Figure 2.**
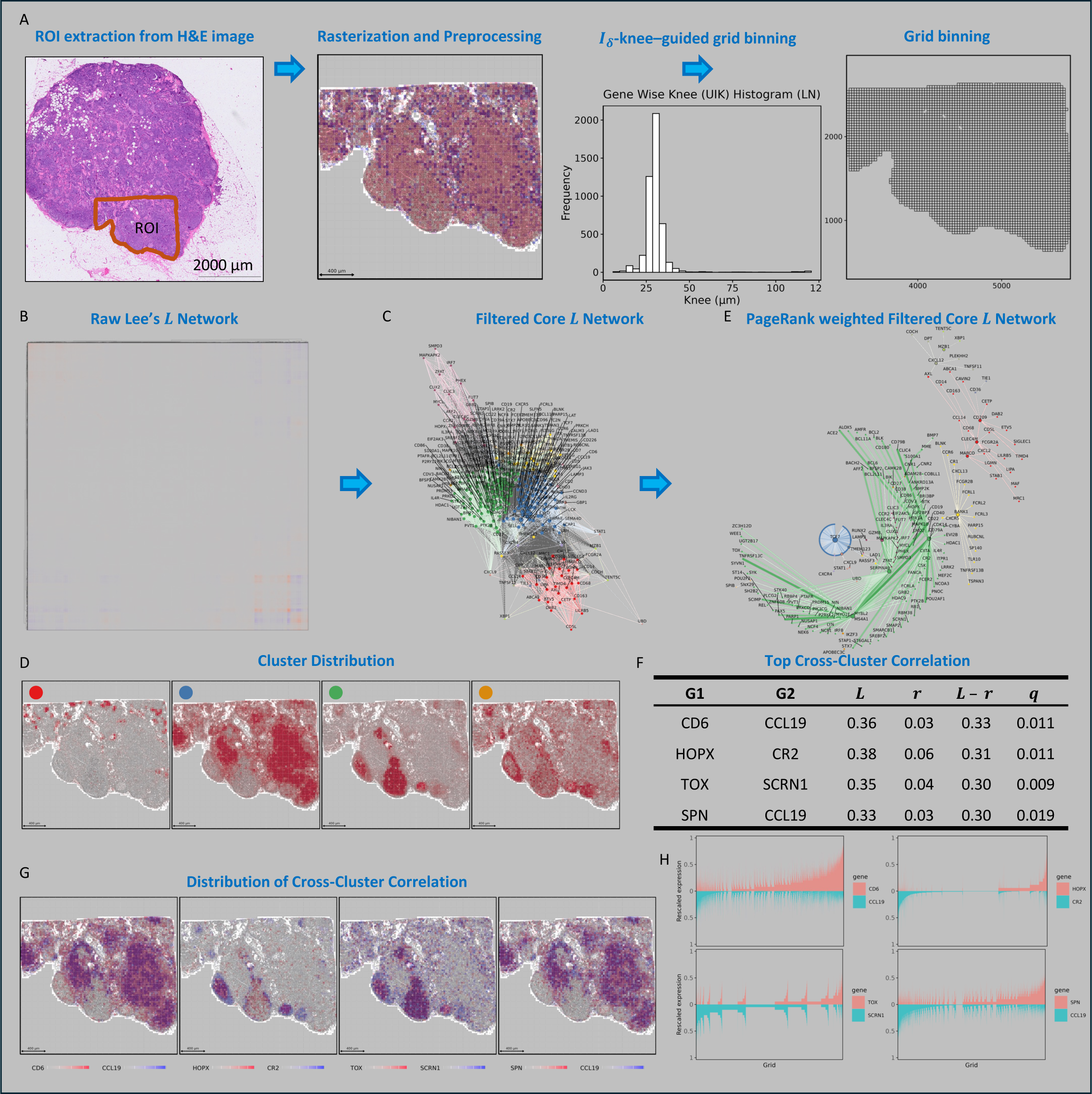
geneSCOPE workflow on 10x Xenium FFPE human LN section (5k Pan-Tissue & Pathways gene panel, ref. [42]) (A) ROI definition and grid binning. An H&E-guided region of interest in a human lymph node section was selected and transcripts were aggregated into 30 μm square grid bins. The mode of histogram of per-gene UIK selects the working width. (B) Raw spatial association. Lee’s *L* was computed for all gene pairs on the normalized grid bin expression matrix to construct the raw Lee’s *L* network. (C) Core network. The spatial network was filtered to retain the top 5% of *L* edges that remained significant at *q* ≤ 0.05, yielding a robust Lee’s *L* core network. (D) Module footprints. Spatial distribution heatmaps for modules 1–4 (of 15 modules, defined on the core network). Each panel shows one module, with bins colored by module’s density; colored dots indicate module identity (1: red, 2: purple, 3: blue, 4: green). (E) Hierarchical relationships. A PageRank-weighted dendro-network (MST backbone; see Methods) summarizes relationships among spatial gene modules. (F) Top *L* − *r* pairs (table). Top four high-*L*, low-*r* gene–gene pairs with *q* ≤ 0.05 (spatially enriched relative to global single-cell co-expression). (G) Spatial maps of pairs. Heatmaps showing the spatial distribution of the same top *L* – *r* pairs across grid bins. (H) Mirror plots. Mirror plots of the grid-wise distributions for these pairs, highlighting neighborhood co-localization versus segregation. Abbreviations: FFPE, Formalin-Fixed Paraffin-Embedded; LN, lymph node.

In the lymph node, comparing spatial versus single-cell gene correlations again revealed biologically meaningful neighborhood signals. The top spatially co-enriched gene pairs recapitulated canonical lymph node microanatomy. For example, CD6–CCL19 and SPN–CCL19 co-localized within paracortical T cell zone territories, whereas HOPX–CR2 (CR2 encodes CD21) was enriched at the T–B interface demarcating the GC perimeter. Likewise, TOX–SCRN1 co-occurred in medullary and sinus regions. These descriptive co-enrichments (often involving non-classical ligand-receptor pairs) mark co-localized gene expression within established anatomical niches, underscoring geneSCOPE’s potential to uncover novel spatial adjacency relationships. Taken together, this lymph node case study demonstrates that leveraging raw molecule coordinates is sufficient to resolve fine-grained intratissue architecture, and that geneSCOPE can distinguish functional subregions across a complex lymphoid tissue while nominating specific adjacent gene pairs for further investigation.

### Resolution Selection and Multiscale Validation

To choose an analysis scale that preserves biological structure while controlling noise, we profiled gene-wise Morisita’s *I*_*δ*_ across grid widths and estimated per-gene UIKs and selected the mode of their distribution. Across three CRC sections (P1, P2, P5) and a human lymph node, the mode of the per-gene UIK distribution consistently fell near 30–35 μm (≈2–3 cell diameters): about 35 μm for CRC P1/P5 and about 31 μm for CRC P2 and lymph node. We therefore used 30 μm as a default and compared against 10 μm (finer) and 55 μm (coarser) resolutions (Fig. 3).

**Figure 3.**
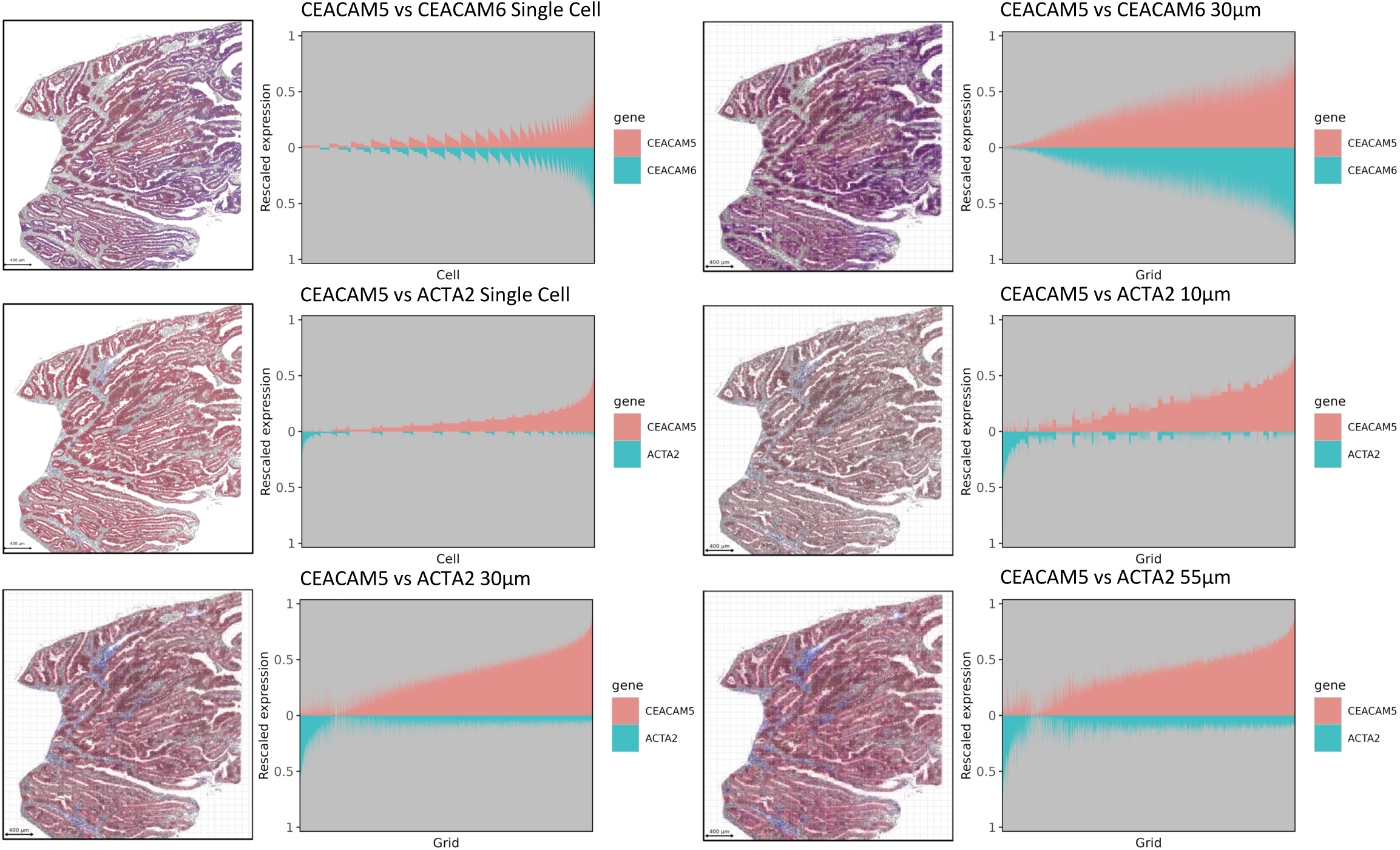
Cell-type markers show minimal cross-bleed across spatial resolutions. A co-localization heatmap of two epithelial tumor markers shows that CEACAM5 (red) and CEACAM6 (blue) occupy nearly identical spatial territories; their overlap appears purple. Across 10 μm, 30 μm, and 55 μm grid binning analyses, markers from different cell types remain spatially segregated—CEACAM5 (red) for cancer epithelium versus ACTA2 (α-SMA; blue) for fibroblasts—indicating minimal cross-bleed between cell-type channels. As a reference, we also show a co-localization heatmap computed from single-cell-level counts with visualization at single-cell centroids.

As a biological check on cross-scale behavior, co-localization heatmaps showed that same-lineage markers remain tightly co-distributed, whereas cross-lineage pairs remain segregated, regardless of bin size. In CRC, CEACAM5–CEACAM6 (epithelial) overlap nearly perfectly, while CEACAM5–ACTA2/α-SMA (epithelium vs fibroblasts) remain spatially distinct at 10, 30, and 55 μm; in lymph node, ITGB2–PTPN6 (pan-leukocyte) co-localize, whereas ITGB2–PDGFRA (immune vs fibroblast) remain separated. Single-cell centroid maps provide a reference view alongside the binned analyses (Fig. 3 and Fig. S4).

Methodologically, Lee’s *L* computed at 10, 30, and 55 μm produced robust association-score distributions suitable for network analysis, and *L* versus single-cell Pearson’s *r* scatterplots retained the expected structure across scales (Fig. S5).

### Spatially Specific Gene–Gene Pairs Revealed by geneSCOPE

Integrating the single-cell reference allowed us to pinpoint gene-gene relationships driven specifically by spatial context. We plotted each gene pair’s spatial co-occurrence (Lee’s *L*) against its global co-expression (Pearson’s *r*) to identify outliers enriched for neighborhood-specific coordination. In the CRC and lymph node dataset, this *L* vs *r* analysis revealed a prominent cluster of gene pairs with weak overall co-expression yet strong local co-localization (Table 1; Fig. 4A; Fig. S6, S7, S8).

**Figure 4.**
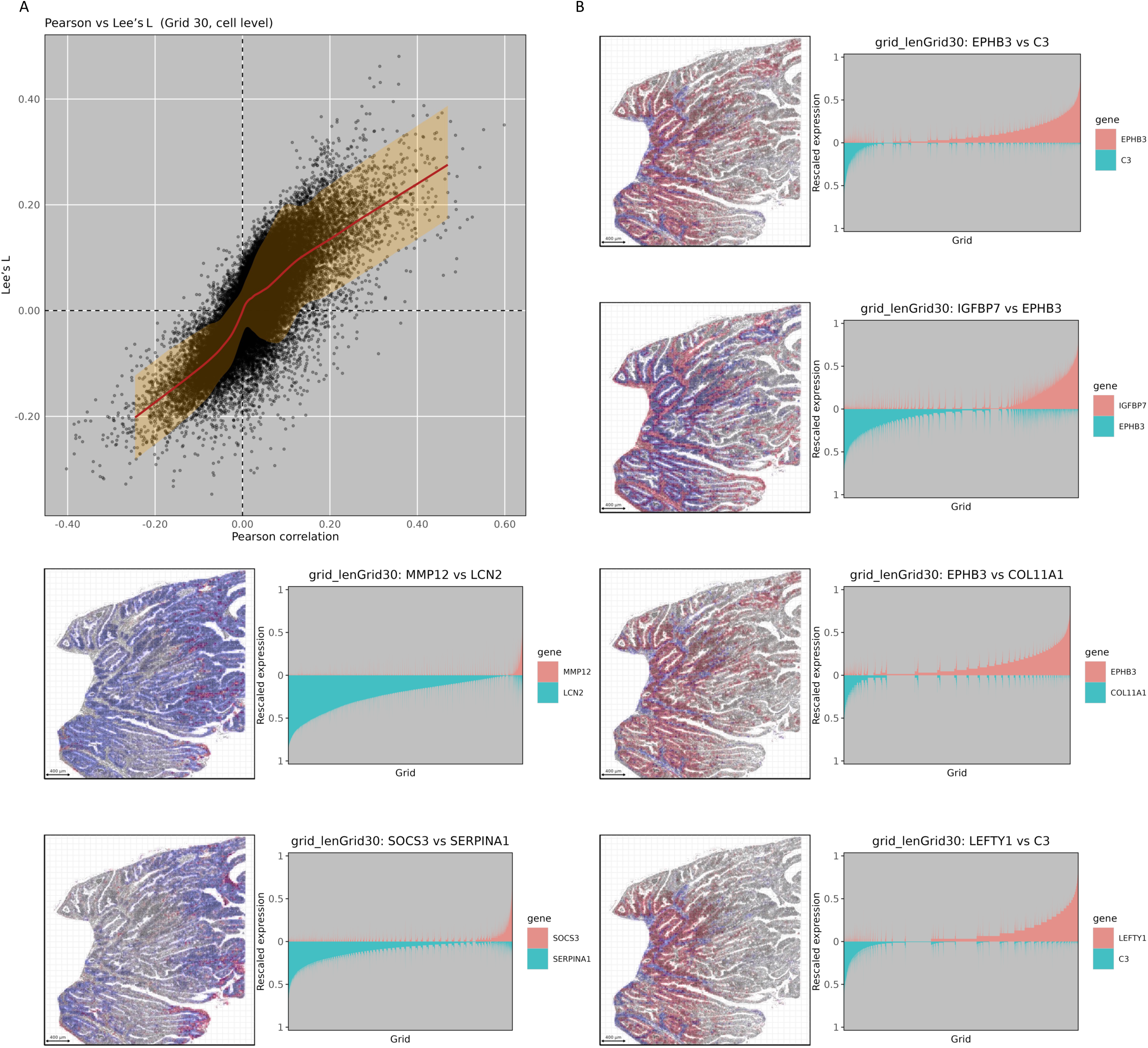
Relationship between spatial Lee’s *L* and Pearson’s *r* in CRC patient P5. (A) Scatter plot of *L* (y-axis) versus single-cell *r* (x-axis) for all unique gene pairs shows many high-*L*/low-*r* outliers, i.e. neighborhood-specific associations that are strong in tissue but weak in dissociated cells. The red curve represents LOESS smoothing with confidence band. (B) Spatial heatmaps of representative high-*L*/low-*r* gene pairs confirm localized co-occurrence in tissue despite weak global co-expression.

**Table 1.**
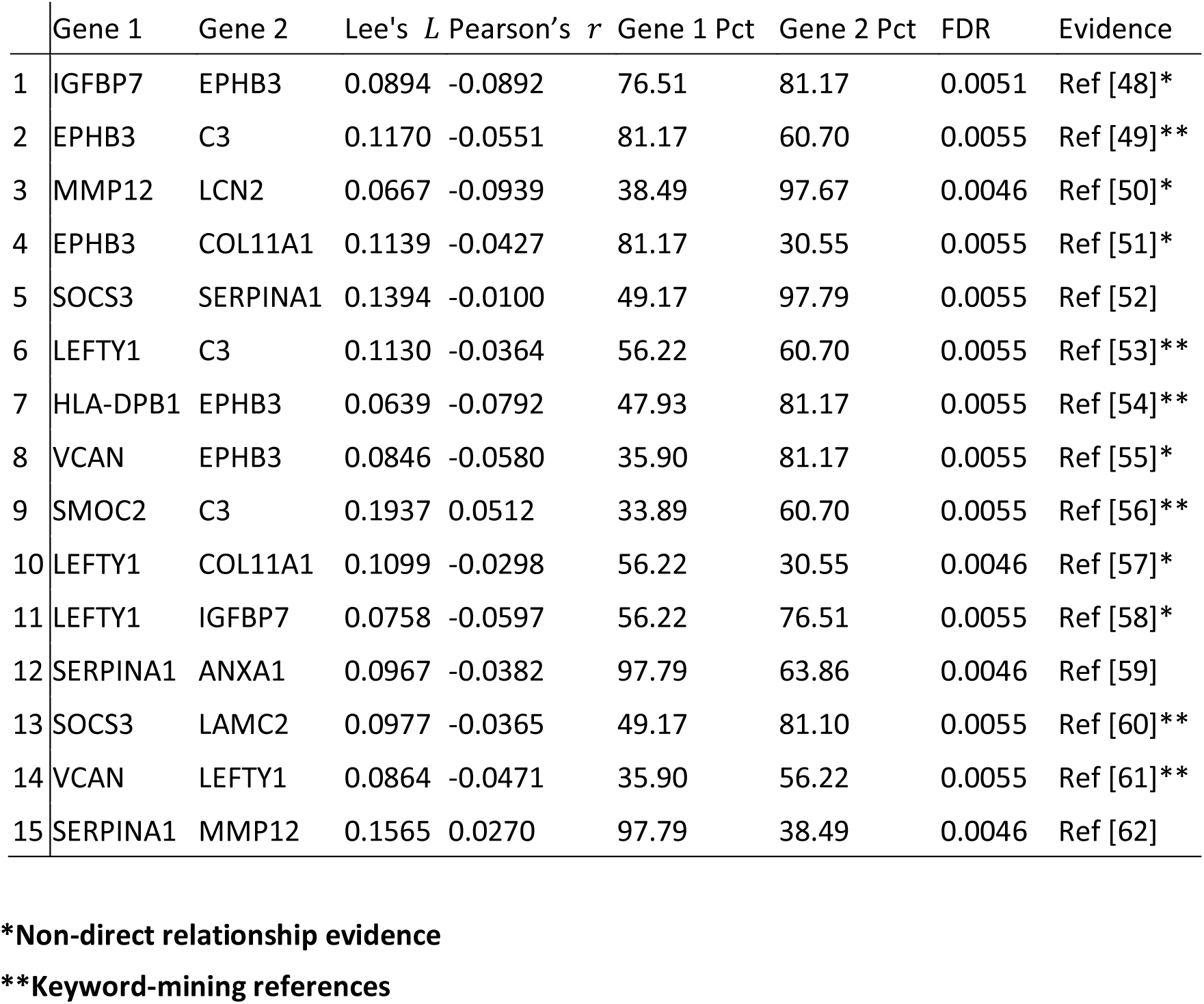
Top 15 gene pairs with high Lee’s *L* (adjacency-weighted spatial association on the 30 μm grid) and low single-cell Pearson’s *r* (global co-expression), identified in the CRC P5 ROI (Xenium). Pairs are ranked by *L* and restricted to those significant after spatially constrained permutations (FDR *q* ≤ 0.05); “Pct” columns indicate the fraction of grid bins with non-zero expression.

| | Gene 1 | Gene 2 | Lee's $L$ | Pearson's $r$ | Gene 1 Pct | Gene 2 Pct | FDR | Evidence |
| --- | --- | --- | --- | --- | --- | --- | --- | --- |
| 1 | IGFBP7 | EPHB3 | 0.0894 | -0.0892 | 76.51 | 81.17 | 0.0051 | Ref [48]* |
| 2 | EPHB3 | C3 | 0.1170 | -0.0551 | 81.17 | 60.70 | 0.0055 | Ref [49]** |
| 3 | MMP12 | LCN2 | 0.0667 | -0.0939 | 38.49 | 97.67 | 0.0046 | Ref [50]* |
| 4 | EPHB3 | COL11A1 | 0.1139 | -0.0427 | 81.17 | 30.55 | 0.0055 | Ref [51]* |
| 5 | SOCS3 | SERPINA1 | 0.1394 | -0.0100 | 49.17 | 97.79 | 0.0055 | Ref [52] |
| 6 | LEFTY1 | C3 | 0.1130 | -0.0364 | 56.22 | 60.70 | 0.0055 | Ref [53]** |
| 7 | HLA-DPB1 | EPHB3 | 0.0639 | -0.0792 | 47.93 | 81.17 | 0.0055 | Ref [54]** |
| 8 | VCAN | EPHB3 | 0.0846 | -0.0580 | 35.90 | 81.17 | 0.0055 | Ref [55]* |
| 9 | SMOC2 | C3 | 0.1937 | 0.0512 | 33.89 | 60.70 | 0.0055 | Ref [56]** |
| 10 | LEFTY1 | COL11A1 | 0.1099 | -0.0298 | 56.22 | 30.55 | 0.0046 | Ref [57]* |
| 11 | LEFTY1 | IGFBP7 | 0.0758 | -0.0597 | 56.22 | 76.51 | 0.0055 | Ref [58]* |
| 12 | SERPINA1 | ANXA1 | 0.0967 | -0.0382 | 97.79 | 63.86 | 0.0046 | Ref [59] |
| 13 | SOCS3 | LAMC2 | 0.0977 | -0.0365 | 49.17 | 81.10 | 0.0055 | Ref [60]** |
| 14 | VCAN | LEFTY1 | 0.0864 | -0.0471 | 35.90 | 56.22 | 0.0055 | Ref [61]** |
| 15 | SERPINA1 | MMP12 | 0.1565 | 0.0270 | 97.79 | 38.49 | 0.0046 | Ref [62] |

\*Non-direct relationship evidence
\*\*Keyword-mining references

Leveraging the high- *L* /low- *r* cluster, geneSCOPE prioritized several candidate neighborhood-conditioned gene–gene pairs at the CRC invasive front. For instance, LEFTY1—a secreted antagonist in the NODAL/TGF-β/SMAD pathway—was co-localized in neighborhoods enriched for cancer-associated fibroblast (CAF) signals, notably complement C3 and IGFBP7 (Fig. 4B). Likewise, EPHB3—an Eph receptor whose down-regulation is associated with increased CRC invasiveness—appeared near C3/IGFBP7-rich stromal regions, consistent with Eph-mediated boundary effects at tumor-stroma interfaces (Fig. 4B). These associations are correlative but biologically plausible: C3⁺ fibroblasts and IGFBP7-secreting stromal cells are implicated in matrix remodeling, immune modulation, and invasiveness, providing a potential mechanistic context for their localized co-occurrence with adjacent epithelial programs. Furthermore, the lymph node analysis benefited from the high gene plexity (about 5,000 genes) of the Xenium platform, which increased the number of measurable gene pairs and likely contributed to the broader spread observed in the *L* vs *r* space. Overall, geneSCOPE’s spatial analysis separates spatially driven co-enrichment from generic co-expression, highlighting neighborhood-level interactions. For example, our results nominate the LEFTY1-C3/IGFBP7 and EPHB3-C3/IGFBP7 pairings, both relatively under-explored in the literature, as candidates for targeted validation *in situ*.

### Spatial Gene Communities and an Illustrative Cross-Niche Path

Using the optimized parameters, we identified 24 spatial gene modules spanning 172/422 genes in the CRC P5 section (Fig. 5A–B). Because Lee’s *L* weights spatial adjacency, module score maps aligned with microanatomy (e.g., invasive front, luminal surface, fibroblast-rich regions) (Fig. 5, S11; Figs. S12-S14 for P1, P2, and LN). To quantify module coherence, we used Jaccard similarity of binarized spatial footprints: across P1, P2, P5 and LN, genes showed systematically higher mean similarity to genes within their own module than to genes in other modules (paired Wilcoxon tests: P1 *p* = 7.13 × 10⁻^29^; P2 *p* = 1.93 × 10⁻^19^; P5 *p* = 4.62 × 10⁻^30^; LN *p* = 1.03 × 10⁻^43^), supporting that the modules capture internally coherent and distinct spatial programs (Fig. S15).

**Figure 5.**
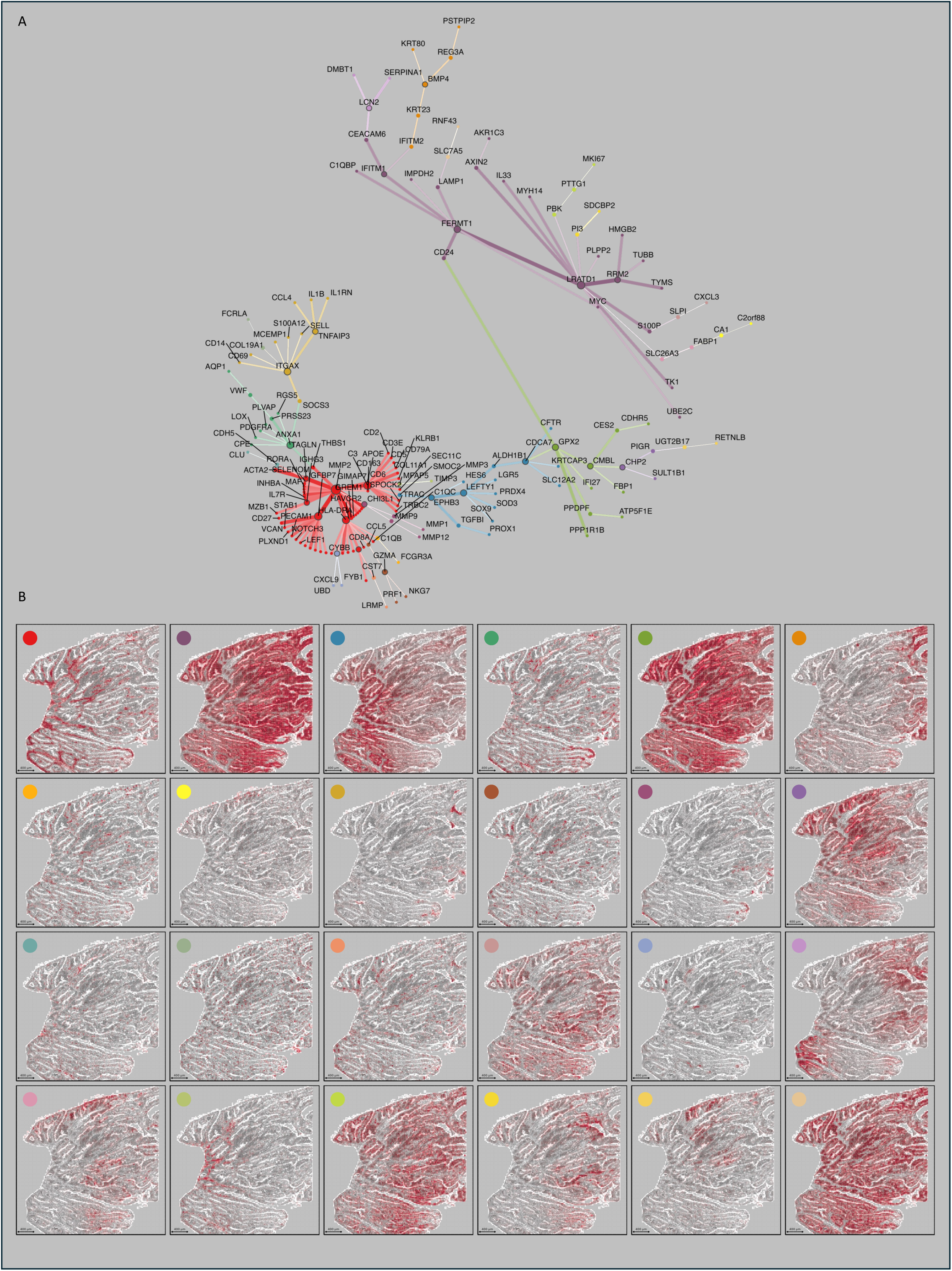
Distinct gene modules occupy discrete spatial niches in CRC patient P5. (A) The core Lee’s *L* network, retaining edges above the 95th percentile of spatial correlation and with FDR-adjusted *q* ≤ 0.05 (log-transformed weights), was clustered by consensus community detection (co-clustering frequency ≥ 0.95) and arranged using a PageRank-weighted dendro-network layout to emphasize hierarchical relationships among the colored gene modules. (B) Heatmaps project each enriched module onto the tissue, with bins colored by the module’s per-bin score and panels keyed to the module’s network color (module colors correspond to network clusters), revealing non-overlapping territories (e.g., invasive margin vs interior epithelium).

In the PageRank-weighted MST representation of the spatial gene network, a stem-like program (module 3; including LGR5 and SMOC2 [39–41]) was positioned between a broadly expressed epithelial program (module 5; including GPX2, PPDPF, and ATP5F1E [42, 43]) and a fibroblast- and ECM-remodeling program (module 1; centered on C3 [44, 45]). This arrangement is consistent with stem-like tumor cells residing at the interface between widely distributed tumor programs and fibroblast-rich, complement- and ECM-associated niches, and illustrates how the backbone layout can help generate hypotheses about the relative spatial positioning of gene programs.

## Discussion

Treating tissues as spatially organized ecosystems reframes microenvironmental influence: putative neighborhood effects become quantities that can be measured directly *in situ*. In keeping with this molecular-based, spatial-association perspective, geneSCOPE takes native molecular coordinates as the primary data, rather than imposing a prior cell taxonomy, and asks how transcriptional programs distribute and co-vary across contiguous neighborhoods of intact tissue. By grounding inference in geography—where proximity shapes phenotype—analyses recover place-specific signals that would be diluted or missed when spatial context is ignored.

Methodologically, the framework distills spatial transcriptomics to its essentials: a grid-based molecular model that aggregates transcripts into spatial bins matched to assay resolution; a spatially weighted correlation (Lee’s *L*) that integrates Pearson’s *r* with spatial weighting to quantify gene-gene co-variation in adjacent regions; and consensus community detection to assemble pairwise associations into coherent spatial modules. This design avoids rigid cell-type aggregation and lets spatial statistics operate directly on *in situ* molecular coordinates, thereby preserving microenvironmental context while sidestepping assumptions carried over from dissociated single-cell workflows. Inevitably, selecting bin size and neighborhood definitions interacts with the modifiable areal unit problem (MAUP) and with threshold choices; to temper these sensitivities, geneSCOPE couples grid tuning to multiscale evaluation so that resolution is chosen where signal and stability balance best—while remaining explicit that analysis parameters influence which patterns are most visible.

Computationally, the same simplifications that respect measurement scale also enable scalability. Binning transcripts and focusing on spatially weighted associations allow geneSCOPE to process sections with about 10^8^ molecules within minutes on a multi-core workstation. In direct runtime benchmarks on the Xenium CRC P5 dataset, geneSCOPE and Hotspot both completed within 10 minutes on the same region of interest and hardware, with comparable wall-clock runtimes (Extended Information). In practice, however, very fine spatial resolutions can reduce power due to sparser counts per bin, whereas coarser resolutions improve stability at the cost of local detail; judicious pre-filtering, edge-weight transformations, and consensus procedures help stabilize networks under these opposing pressures. Panel breadth and molecule density therefore remain practical determinants of detectable effect size: low capture or narrow panels can obscure subtle neighborhood signals, while high-plex assays broaden the accessible space of spatial associations.

Across tissues, this approach reveals molecular patterns alongside known anatomy and is especially sensitive to boundaries and interfaces. Without pre-annotating cell types, geneSCOPE recapitulates canonical microanatomy—such as germinal-center (GC) and peri-GC territories in human lymph node—and nominates localized, context-dependent co-occurrences at tissue interfaces (e.g., HOPX-CR2 at the T-B border) that suggest specialized adjacency programs worthy of follow-up. More broadly, the framework reveals immune-epithelial/stromal neighborhood-level associations that help rationalize co-varying phenomena such as immune infiltration, stromal remodeling, and epithelial plasticity—consistent with ecological metrics in oncology where immune-tumor colocalization carries prognostic value. These module-level gradients also resonate with developmental atlases in which gene expression varies continuously along tissue axes rather than as step functions. While these observations are inherently correlational, they convert spatial adjacency into a compact set of testable hypotheses for targeted *in situ* validation and perturbation.

Generalizability emerges from the same measurement-aware stance. Across colorectal tumor and lymphoid tissue, an effective neighborhood scale on the order of 30-35 μm—roughly two to three cell diameters—proved sufficient to capture short-range coordination while maintaining stability, offering a pragmatic default with clear biological interpretation. Because platforms span complementary resolution-throughput regimes (multi-cell spots in Visium versus single-molecule imaging in MERFISH/Xenium), geneSCOPE adapts by parameterizing its grid and neighborhood definitions; at 10, 30, or 55 μm, it recovers robust association distributions suitable for network analysis, with finer scales prioritizing detail and coarser scales improving power and noise tolerance. Inevitably, however, 2D sections provide incomplete views of 3D tissue architecture. Multiscale binning, ROI-aware null models, and cross-section consistency checks can reduce 3D sampling biases, but cannot eliminate them; extending analyses to registered serial sections and volumetric spatial omics will further mitigate these limitations as datasets mature [9, 10, 46].

Integration with complementary modalities helps separate spatially driven coordination from general co-expression. Treating single-cell RNA-seq as a non-spatial baseline and contrasting Pearson’s *r* with *in situ* Lee’s *L* highlights gene pairs whose concordance is conditioned by neighborhood context rather than by cell-intrinsic programs, a pattern seen in both tumor and lymphoid tissues. Likewise, overlaying spatial proteomics and morphology enables triangulation of transcript modules against protein abundance and structural cues, as recent cardiovascular and hematologic multimodal studies illustrate [6, 47]. These comparisons sharpen interpretation but are not themselves evidence of causality; moving from spatial association to mechanism ultimately requires targeted, spatially resolved perturbations, ideally designed around the module interfaces that geneSCOPE brings into focus.

In summary, geneSCOPE extracts neighborhood-level co-expression with minimal assumptions, embeds those programs back onto the tissue map, and turns spatial adjacency into testable, mechanism-oriented hypotheses. The framework is portable across platforms, tissues, and modalities; it is also explicit about scale choices, panel and density constraints, and the unavoidable 2D-to-3D sampling biases that it can attenuate but not abolish. As spatial multi-omics and 3D atlases expand, we anticipate geneSCOPE will help systematize “module-interface-pathway” interaction maps within complex tissues, advancing mechanistic understanding and nominating spatially informed targets for precise intervention.

## Acknowledgements

This work was supported by the Otsuka Toshimi Scholarship Foundation. We used ChatGPT for English editing/translation and coding assistance. These tools did not generate figures, references, or substantive scientific content, and no AI system is listed as an author. All AI-assisted outputs were reviewed and verified by the authors, who take full responsibility for the content, in line with the COPE position statement on Authorship and AI.

## Key Points

We developed geneSCOPE, a tool that identifies spatial gene modules—sets of genes with coordinated, location-dependent expression.

The geneSCOPE pipeline also detects gene pairs that exhibit spatially coordinated co-expression across different cell types, revealing neighborhood-conditioned interactions.

We propose a data-driven grid-size selection method—using the mode of the per-gene UIK distribution derived from Morisita’s *I*_*δ*_–width curves—to leverage spatial information in gene distributions.

## Author contributions

**S. Zhang:** Methodology, Software, Data Curation, Investigation, Formal analysis, Visualization, Writing. **K. Saeki:** Methodology, Formal analysis, Investigation, Writing. **H. Haeno:** Conceptualization, Methodology, Data Curation, Investigation, Formal analysis, Supervision, Writing.

## Short description of the authors

Shicheng Zhang is a PhD student in the graduate school of biological sciences at Tokyo University of Science.

Koichi Saeki is an assistant professor in the research institute for biomedical sciences at Tokyo University of Science.

Hiroshi Haeno is an associate professor in the research institute for biomedical sciences at Tokyo University of Science.

---

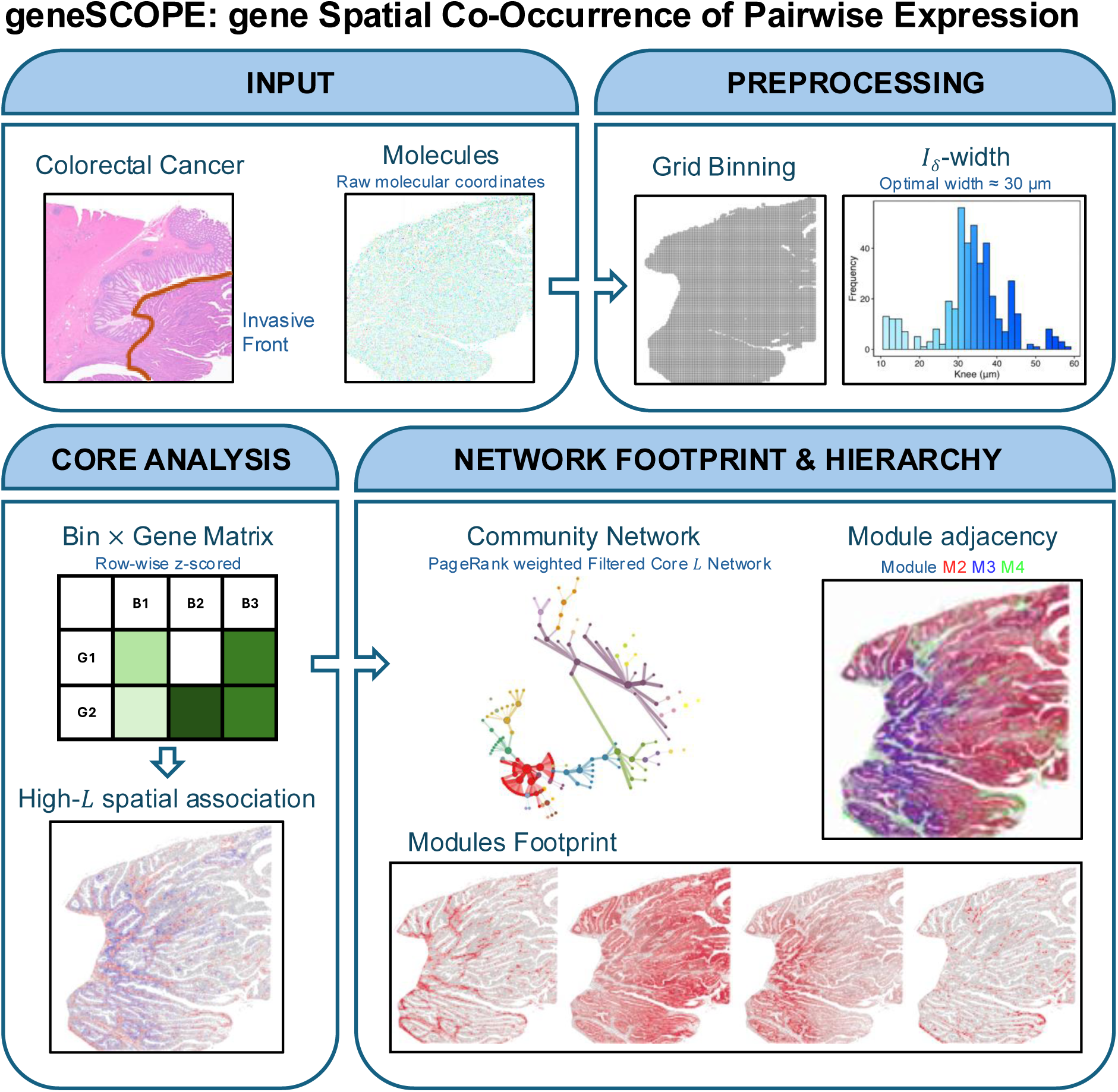

**Figure S1.**
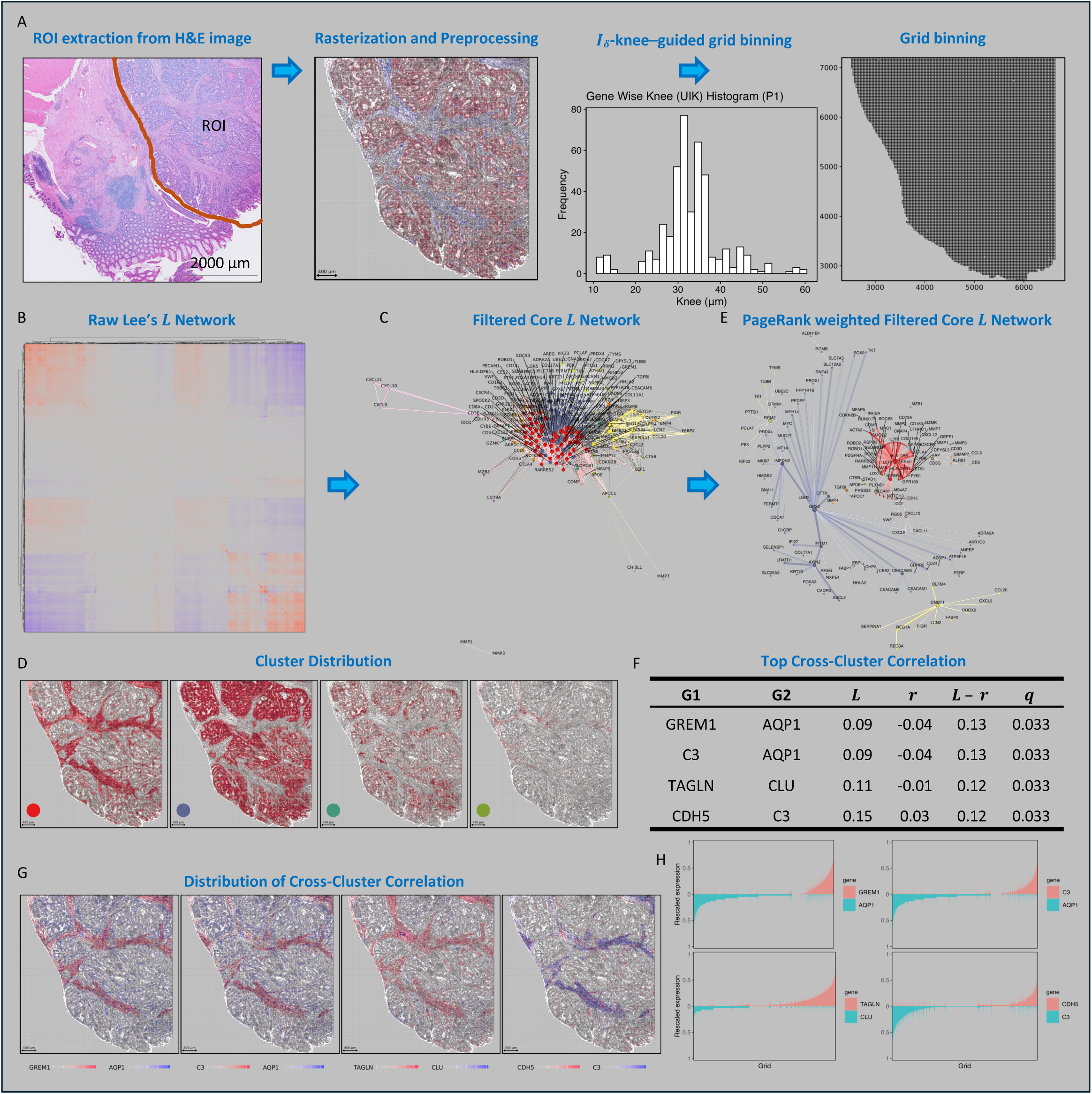
geneSCOPE workflow on CRC patient P1 (GSE280314).

