## Supplemental Information for "geneSCOPE: gene Spatial Co-Occurrence of Pairwise Expression"

**Extend Information**

**Benchmarking: hotspot Pipeline (Python)**

**Overview:** We employed a custom Python script implementing the hotspot algorithm for spatial gene module discovery. The hotspot is a graph-based procedure originally developed for single-cell genomics that identifies informative genes and clusters them into co-expression modules based on spatial autocorrelation patterns. By running this pipeline on our data, we obtained a benchmark for comparison against geneSCOPE's results under identical data and region conditions.

**Input Data and ROI:** The hotspot pipeline accepts spatial transcriptomics data either in a legacy matrix format (with counts.mtx, genes.tsv, and coords.tsv files) or in 10x Genomics Xenium output format (HDF5/MTX count matrix plus a cells.csv coordinates file). In both cases, it ingests a gene expression count matrix along with each cell's spatial coordinates. An optional polygon ROI (region of interest) CSV can be provided to confine the analysis to a specific tissue area. For consistency, we used the same fixed ROI for all runs, ensuring that hotspot analyzed the identical subset of cells/molecules as geneSCOPE. This ROI restriction means hotspot's clustering was performed on the exact same tissue region used in the geneSCOPE analysis.

**Spatial Graph Construction:** After loading the data, the pipeline builds a k-nearest neighbor (KNN) graph connecting each cell to its 15 nearest neighboring cells in physical space. This graph captures local spatial proximity relationships and forms the basis for downstream autocorrelation calculations. We fixed the neighborhood size at k = 15 for all analyses.

**Spatial Autocorrelation Analysis:** Next, hotspot computes a spatial autocorrelation statistic for each gene across the neighbor graph. This per-gene statistic quantifies how much that gene's expression is spatially clustered or dispersed across the tissue. Genes with high positive autocorrelation (i.e., strongly clustered expression) are labeled as "informative" spatial genes and are carried forward for module detection, while genes with near-random or over-dispersed patterns are not considered further. This gene filtering step focuses the analysis on spatially patterned genes.

**Gene-Gene Local Correlation and Clustering:** The pipeline then calculates pairwise local correlation between the expression profiles of all informative genes across neighboring cells. This step identifies genes that show similar spatial expression patterns. The hotspot leverages the shared information among nearby cells to improve sensitivity in detecting these gene-gene relationships. The hotspot performs hierarchical clustering on the gene-gene local correlation matrix to group genes into modules of spatially co-expressed genes. This yields gene modules, each representing a set of genes with strongly coordinated spatial expression. The hotspot model was run with its default distribution (depth-adjusted negative binomial, appropriate for UMI count data) to maintain consistency.

**Outputs:** The hotspot script outputs a per-gene spatial autocorrelation score for every gene, along with an associated significance (FDR) and the gene module assignments for informative genes. We used these results, the list of spatially patterned genes and their module groupings, to compare and evaluate overlap with geneSCOPE's gene modules.

**Runtime Benchmarking:** To measure performance, we executed the hotspot analysis via a dedicated benchmarking shell script for each dataset. This wrapper script launches the hotspot pipeline while monitoring resource usage and timing. It logs the runtime for each sample (along with CPU metrics) to a file for reproducibility. Importantly, this monitoring utility does not alter the hotspot analysis itself; it only measures how long each run takes. All runs were processed on the same hardware, using 16 parallel threads and 32 GB memory, ensuring a consistent environment to compare runtimes across runs and against geneSCOPE's execution times.

**Benchmarking: geneSCOPE Pipeline (R)**

**Overview:** We carried out this part of the study using a core R script that implements the geneSCOPE pipeline. This R script constitutes the main computational framework for geneSCOPE and was used for the key analyses reported in the manuscript. It embodies the full pipeline logic, from input data handling to network clustering, focusing on the calculation of spatial gene co-expression metrics and the identification of gene modules. By running geneSCOPE on the same data and region as hotspot, we directly compare their performance and results.

**Input Data and Grid Binning:** The geneSCOPE pipeline begins with a gene-by-bin count matrix derived from spatially gridded transcript data. The pipeline can accept raw 10x Genomics Xenium outputs as input and then aggregate counts into a spatial grid. All detected transcripts within the defined ROI (identical to the one used in hotspot script) are binned into fixed-size spatial units (30 µm grid size in this analysis) to produce the input matrix. Each bin thus represents a small tissue neighborhood, providing a balance between spatial resolution and noise reduction.

**Spatial Weight Matrix Construction:** After binning, the pipeline constructs a spatial weights matrix that captures the adjacency relationships among the bins. Using the regular grid layout, each bin is considered adjacent (a neighbor) to those that share either a boundary or a corner with it (the queen contiguity rule). This results in a lattice-like neighborhood graph of bins across the tissue. The spatial weights matrix is typically binary (1 for neighbors, 0 for non-neighbors), and it defines which bin pairs are considered "neighbors" for subsequent calculations.

**Spatial Co-expression Calculation (Lee's L):** Using the binned expression matrix and the spatial weights matrix, the script computes Lee's L for every pair of genes. Lee's L is a bivariate spatial correlation statistic that quantifies the degree to which two genes' expression levels co-vary across neighboring spatial bins - effectively measuring spatial co-expression or co-enrichment. A positive L indicates that two genes tend to be high (and low) together in adjacent bins more often than expected by chance (suggesting they are co-expressed in the same locales), whereas an L near zero suggests little to no spatial association between the two genes. This comprehensive pairwise analysis yields a gene-gene spatial association matrix (with L values as edge weights) that underpins the gene network. Notably, because Lee's L inherently accounts for the spatial structure (using the defined neighbors), genes that are mapped to the same tissue niches or structures will show high L values.

**Clustering of Spatially Co-expressed Genes:** The resulting gene-gene association matrix is then analyzed to detect clusters (modules) of spatially co-expressed genes. We treat the genes as nodes in a network, with edges weighted by their L correlation values, and apply a consensus clustering approach based on the Leiden community algorithm to identify groups of genes with similar spatial expression patterns. The outcome is a set of gene modules, each representing a group of genes that co-localize in the tissue and share a distinct spatial expression profile across the 30 µm bins. Because Lee's L accounts for local proximity, genes grouped into the same module correspond to coherent tissue domains at approximately the 2-3 cell diameter scale. These modules of spatially co-expressed genes form the basis for downstream biological interpretation, such as identifying cell types or tissue structures associated with each module.

**Outputs:** The results include the gene-gene L matrix, containing the spatial co-expression score for each gene pair, associated significance (FDR) for the L values, and a list of gene modules and their constituent genes. Each module is essentially a cluster of genes with high mutual L values.

**Runtime Benchmarking:** We processed all runs with the geneSCOPE R pipeline using a consistent set of parameters and computing resources. The spatial bin size was fixed at 30 µm for all analyses to maintain a uniform scale across runs. The script was executed with 16 parallel threads and 32 GB memory for each run. For reproducibility and timing, we employed a simple wrapper shell script to launch the R pipeline for each dataset and record its execution time. By running geneSCOPE on the same hardware and threads as hotspot, we were able to compare runtimes directly in a consistent manner.

**Benchmark Results and Clustering Comparison**

For performance benchmarking, we executed hotspot and geneSCOPE five times on the P5 dataset using the same ROI as in the manuscript. Across runs including the most extreme observed run, both methods completed within 10 minutes. hotspot was generally faster (most runs finished in about 300 s), whereas geneSCOPE typically completed in about 400 s.

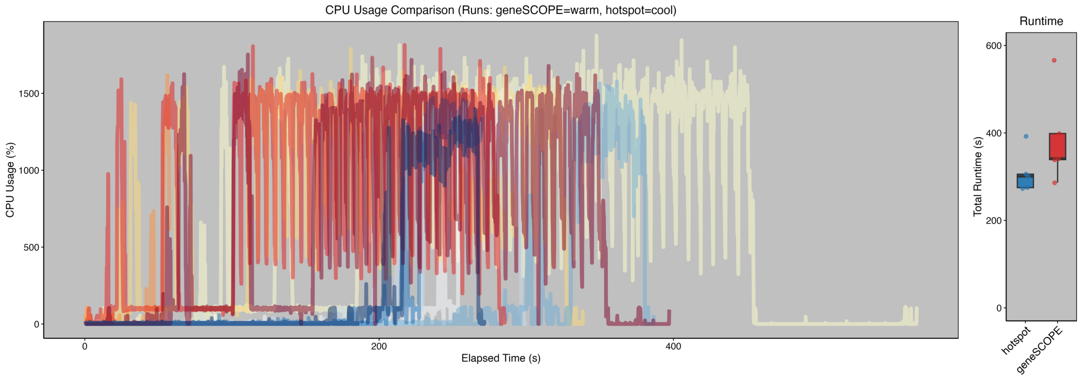

Despite the slightly longer time, geneSCOPE's grid-based binning yields a substantially lower computational burden than a full single-cell Lee's L computation would entail; modest reductions in spatial resolution markedly reduce computational cost while maintaining an effective signal-to-noise ratio.

In terms of clustering outcomes, both geneSCOPE and hotspot captured the overall tissue architecture: delineating features such as the tumor parenchyma (core), stromal band, and vascular network. However, the two methods make different trade-offs between cellular purity and interpretability. The hotspot prioritizes continuity in microniches, often merging co-localized multi-lineage cells at the invasive front or the tumor-stroma interface into one cluster. This yields clusters that are spatially contiguous but contain mixed cell types (lower cell-type purity), which weakens the clarity of marker-based interpretations. In contrast, geneSCOPE produced "cleaner" clusters in this

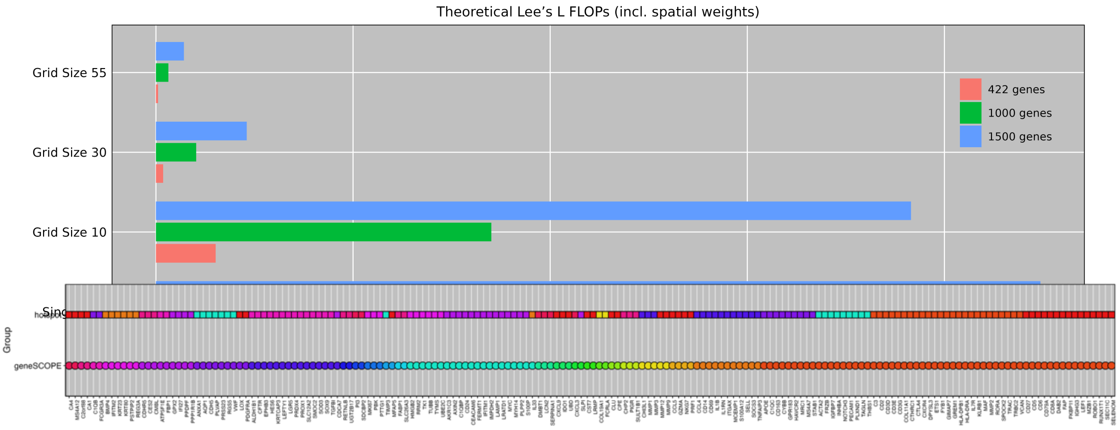


sample: endothelial cells, myeloid cells, proliferative epithelial cells, and differentiated epithelial cells each formed distinct, high-purity clusters with consistent marker profiles. Even at tissue boundaries, geneSCOPE clearly separated epithelial cells, CAFs (cancer-associated fibroblasts), and myeloid cells. This separation enables more intuitive biological interpretation, more robust differential/pathway/regulatory analyses, and stronger comparability and reusability across tissue sections and cohorts. Therefore, in a direct side-by-side comparison on the same data, if one prioritizes cell-type purity, coherence of marker expression in biological processes, and analytical robustness, geneSCOPE holds a clear advantage. Conversely, integrating Hotspot can complement these results by highlighting histological continuity within very small multicellular niches.
